# Molecular Imaging of Collagen Turnover in Myocardial Infarction

**DOI:** 10.64898/2026.01.16.699986

**Authors:** Afarin Neishabouri, Mean Ghim, Onur Varli, Azmi A. Ahmad, Gunjan Kukreja, Zhengxing Zhang, Jie Li, Jakub Toczek, Mani Salarian, Jiasheng Zhang, Daniel Ein Alshaeba, Fadi G. Akar, Chi Liu, S. Michael Yu, Mehran M. Sadeghi

**Author notes:** Equal contribution (Afarin Neishabouri, Mean Ghim). **Corresponding Author**: Mehran M. Sadeghi, M.D., Professor of Medicine (Cardiology), Yale School of Medicine, 300 George Street, Room 770G, New Haven, CT 06511, USA. **First Authors:** Afarin Neishabouri, MD Postdoctoral Associate Mean Ghim, PhD Associate Research Scientist Yale School of Medicine 300 George Street, R2320 New Haven, CT 06511, USA.

## Abstract

Cardiac fibrosis is a key contributor to cardiomyopathy after myocardial infarction (MI). Existing imaging techniques can detect established fibrotic changes; however, they lack sensitivity for ongoing collagen turnover—a dynamic process involving the denaturation of collagen triple helix. Molecular imaging of this process could enhance risk assessment and aid in the development of anti-fibrotic treatments. This study aimed to evaluate ^99m^Tc-(HE)₃-(GPO)₉, a radiotracer designed to target denatured collagen, as a biomarker of collagen turnover after MI.

**Methods:** ^99m^Tc-(HE)₃-(GPO)₉ incorporates glycine-proline-hydroxyproline (GPO) repeats and can hybridize with denatured single- or double-stranded collagen. MI was induced in mice by ligation of the left anterior descending artery; sham-operated animals served as controls. At 2 weeks post-MI, animals underwent myocardial perfusion imaging or contrast-enhanced CT to detect the infarct zone, followed by SPECT/CT imaging using ^99m^Tc-(HE)₃-(GPO)₉ or a control scrambled tracer. Tracer uptake was quantified in vivo and ex vivo with gamma counting and autoradiography. Different aspects of fibrosis were examined by tissue analysis, along with autoradiography with a matrix metalloproteinase-targeted radiotracer, ^99m^Tc-RYM1. Tracer binding was also assessed in human cardiac tissue through ex vivo autoradiography.

**Results:** ^99m^Tc-(HE)₃-(GPO)₉ SPECT/CT revealed significantly higher tracer uptake in the infarct zone of MI mice compared to the remote zone and sham controls (P < 0.0001 for both). Tracer uptake was confirmed by autoradiography, which showed a strong correlation between SPECT and autoradiography (R = 0.81, P < 0.01). The scrambled tracer exhibited minimal cardiac uptake, demonstrating the specificity of ^99m^Tc-(HE)₃-(GPO)₉ signal. Denatured collagen staining and ^99m^Tc-RYM1 autoradiography showed similar patterns as ex vivo ^99m^Tc-(HE)₃-(GPO)₉ autoradiography, while the ratio of denatured collagen to procollagen in the infarct zone significantly increased from day 3 to 2 weeks post-MI. Finally, ^99m^Tc-(HE)₃-(GPO)₉ bound to human fibrotic (but not normal) cardiac tissue.

**Conclusion:** ^99m^Tc-(HE)₃-(GPO)₉ enables non-invasive detection of denatured collagen as a marker of collagen remodeling in vivo, offering a promising tool for assessing fibrotic remodeling after MI. Collagen, procollagen, and denatured collagen, along with MMP activation, exhibit distinct patterns, and their combined imaging may provide a comprehensive molecular fingerprint of cardiac fibrosis, advancing personalized management of cardiomyopathy.

## Introduction

Dysregulated tissue repair in response to injury results in fibrosis, a chronic, progressive process characterized by excessive deposition of extracellular matrix (ECM) proteins, including collagen (*1*). Fibrosis can lead to organ dysfunction, and despite being a major global healthcare burden, the translation of known pathogenic mechanisms into effective antifibrotic therapies has remained challenging. The detection of the processes involved in fibrosis and tracking its progression and regression can accelerate the development and evaluation of new antifibrotic therapies and help stratify patients for targeted personalized antifibrotic treatments (*2*).

Myocardial fibrosis is associated with a wide range of cardiac pathologies, including ischemic and non-ischemic cardiomyopathies (*3*). Ischemic events, such as myocardial infarction (MI), result in replacement fibrosis, the recruitment of inflammatory cells, and the activation of ECM-producing myofibroblasts. Although the resulting collagen-based scar tissue protects the infarcted ventricle from catastrophic mechanical complications such as cardiac rupture, it may contribute to ventricular remodeling, especially when associated with interstitial fibrosis in the remote zone, and lead to systolic and diastolic dysfunction and potentially heart failure (HF) (*4*). HF following acute MI is the most potent predictor of related mortality (*5*).

Non-invasive imaging can detect cardiac fibrosis and its consequences, such as changes in ventricular function and geometry, primarily using echocardiography and cardiac magnetic resonance (CMR) (*6,7*). Myocardial perfusion imaging (MPI), using positron emission tomography (PET) or single-photon emission computed tomography (SPECT), detects the perfusion deficit associated with scar (*8*). Newly introduced molecular imaging approaches can evaluate the established fibrosis and fibrosis-associated processes by targeting integrins, collagen, fibroblast activation protein (FAP), or proteolytic enzymes (*9–12*). While targeting FAP, integrin activation, or matrix metalloproteinases (MMPs) addresses important fibrosis-associated processes (*13–15*), none of these approaches directly detects the changes in collagen structure that mediate collagen turnover.

Collagen molecules have a unique triple-helical structure that unfolds following enzymatic cleavage (e.g., by MMP, cathepsin K) or mechanical damage. Collagen hybridizing peptides (CHPs) are a family of synthetic peptides with repetitive collagen-mimetic sequences, such as glycine–proline–hydroxyproline [(GPO)_n_]. CHPs can hybridize to denatured, single- or double-stranded collagen chains by forming a triple helix, enabling detection of collagen remodeling during fibrosis (*16–18*). We have recently introduced a CHP-based radiotracer, ^99m^Tc-(HE)_3_-(GPO)_9_, that contains a C-terminal (GPO)_9_ as the targeting moiety (*19*), to detect denatured collagen by SPECT/CT imaging. In the current study, we investigated the performance of ^99m^Tc-(HE)_3_-(GPO)_9_ for molecular imaging of collagen turnover in murine MI. In vivo SPECT/CT imaging and ex vivo analysis of tracer uptake revealed higher tracer uptake in the infarct zone, which correlated with the presence of denatured collagen. The spatial and temporal patterns of denatured collagen in relation to other molecular markers of tissue fibrosis were characterized, and the ability of ^99m^Tc-(HE)_3_-(GPO)_9_ to bind to human cardiac fibrosis was demonstrated as a prelude to future clinical translation.

## Materials and Methods

Detailed materials and methods are provided in the supplemental appendix.

### Radiotracers

The (HE)_3_-(GPO)_9_ precursor was synthesized and radiolabeled to generate^99m^Tc-(HE)_3_-(GPO)_9_ as described (*19*). This tracer, which targets denatured collagen, features a polyhistidine–glutamic acid [(HE)_3_] N-terminal sequence for site-specific radiolabeling, linked to a C-terminal targeting moiety consisting of 9 GPO repeats [(GPO)_9_] via a flexible 3-glycine linker. A control tracer, ^99m^Tc-(HE)_3_-(GPO)_SCR_, where (GPO)_9_ is replaced with a scrambled sequence (PGOGPGPOPOGOGOPPGOOPGGOOPPG), was used as a negative control agent.

### Animals

Animal experiments were performed under protocols approved by the Institutional Animal Use and Care Committees of Yale University and the Veterans Affairs Connecticut Healthcare System. To induce MI, 10- to 16-week-old C57BL/6J mice (n = 39, Supplemental Fig. 1) of both sexes underwent left anterior descending (LAD) artery ligation 2 mm distal to the left auricle, as described with some modifications (*20*). Sham-operated or unoperated animals (n = 8, Supplemental Fig. 1) were used as controls.

## Results

### Radiochemistry

^99m^Tc-(HE)_3_-(GPO)_9_ and its control, non-binding tracer, ^99m^Tc-(HE)_3_-(GPO)_SCR_, were radiolabeled with ^99m^Tc-tricarbonyl and purified to a specific activity of 83.9±35.5 GBq/µmol and 88.1±27.2 GBq/µmol, respectively, as described, and their labeling efficiency and purity were confirmed by radio-HPLC (Supplemental Fig. 2).

### ^99m^Tc-(HE)_3_-(GPO)_9_ SPECT/CT in MI

Echocardiography at 2 weeks after LAD ligation showed the presence of regional wall motion abnormalities and a significant reduction in LVEF, as compared to sham-operated animals (P < 0.01, Supplemental Fig. 3). ^99m^Tc-tetrofosmin myocardial perfusion imaging or eXIA 160XL CE-CT imaging at 2 weeks after surgery confirmed the presence of an anterolateral infarct area in MI animals, while, as expected, sham-operated animals did not show any perfusion defect (Fig. 1A). Myocardial perfusion imaging was followed by ^99m^Tc-(HE)_3_-(GPO)_9_ microSPECT/CE-CT imaging 2-3 days apart, starting at 60 min after tracer injection. ^99m^Tc-(HE)_3_-(GPO)_9_ SPECT/CT images showed markedly higher uptake of the tracer in the infarct zone in MI-induced animals compared to their remote zone and compared to sham-operated mice (Fig. 1A). This was confirmed by quantifying the LV wall ^99m^Tc-(HE)_3_-(GPO)_9_ signal on SPECT/CT images [0.65 %ID/mL (0.56–0.76 %ID/mL) for the infarct zone versus 0.46 %ID/mL (0.37–0.54 %ID/mL) for the remote zone in the MI group (n = 18, P < 0.0001), and 0.34 %ID/mL (0.31–0.39 %ID/mL, n = 5) in the corresponding anterolateral wall of sham-operated animals, P < 0.0001]. There was no difference in tracer uptake between the anterolateral wall and the posteroseptal wall of sham-operated animals (Fig. 1B).

**Figure 1:**
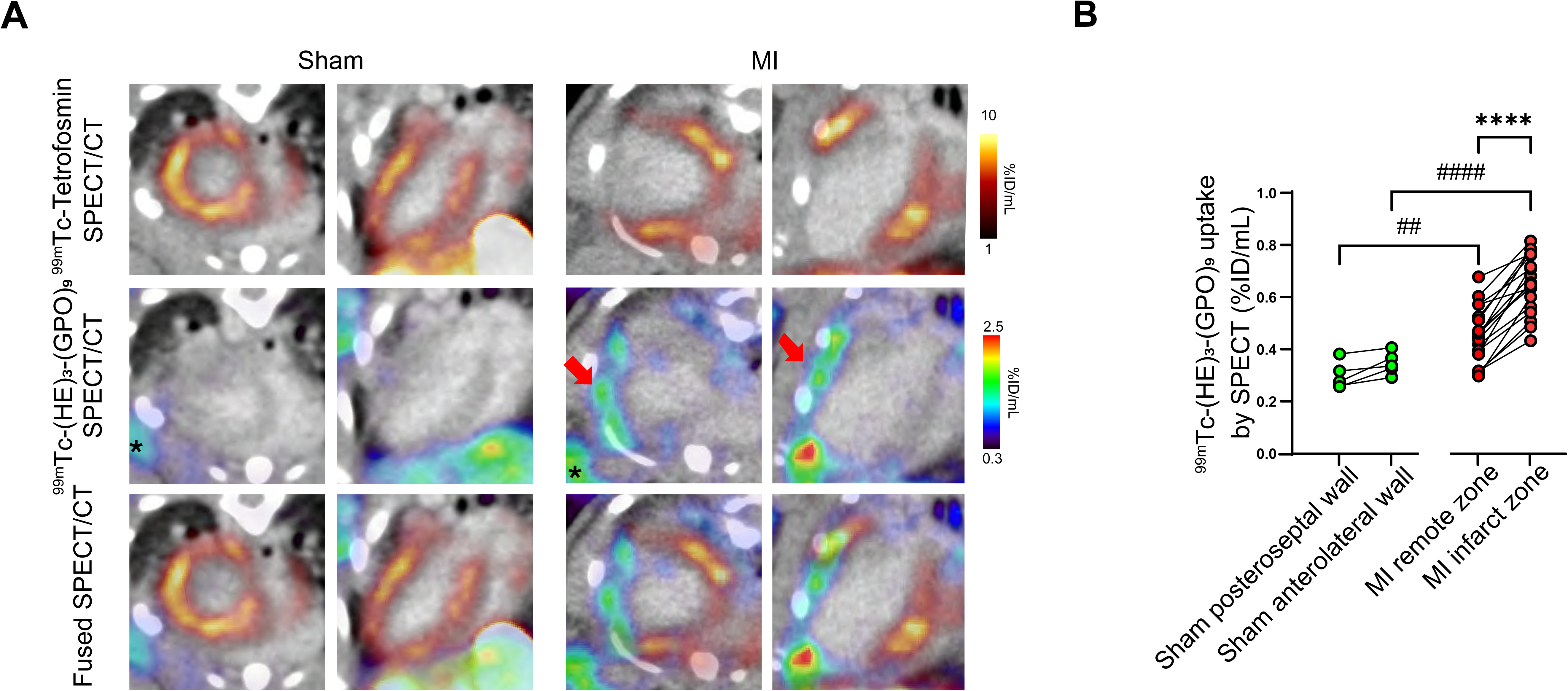
^99m^Tc-(HE)_3_-(GPO)_9_ SPECT/CT in myocardial infarction. (A) Illustrative examples of in vivo SPECT/Exitron Nano-12000-enhanced CT images of sham-operated and MI-induced mice at two weeks after surgery. Red arrows in the middle row indicate ^99m^Tc-(HE)_3_-(GPO)_9_ uptake in the infarct zone of MI animals, identified by ^99m^Tc-tetrofosmin myocardial perfusion imaging (top row). The lower row shows fused ^99m^Tc-tetrofosmin and ^99m^Tc-(HE)_3_-(GPO)_9_ images. The stars mark ^99m^Tc-(HE)_3_-(GPO)_9_ uptake at the surgical site in both sham and MI animals. (**B**) Quantification of the ^99m^Tc-(HE)_3_-(GPO)_9_ SPECT signal in the infarct and remote zones of MI-, and the corresponding walls of sham-operated mice. ID: injected dose. ##: P < 0.01, ####: P < 0.0001, Mann-Whitney *U* test; ****: P < 0.0001, Wilcoxon test.

### ^99m^Tc-(HE)_3_-(GPO)_9_ biodistribution

Ex vivo evaluation by gamma-well counting, following animal euthanasia at 2 hours, showed significantly higher tracer uptake in the apex of MI mice than in the sham group (1.31 vs. 0.46 %ID/g, P = 0.01), with no difference in residual blood radioactivity. Furthermore, tracer uptake in the lung, liver, spleen, kidney, white adipose tissue, and muscle was not significantly different between the sham and the MI groups (Supplemental Fig. 4A). Importantly, there was a significant correlation between the SPECT-based (in %ID/mL) quantification of tracer uptake in the apex and its ex vivo, gamma-well counting–based (in %ID/g) quantification of tracer uptake in the same animals (Spearman’s R = 0.71, P < 0.05; Supplemental Fig. 4B), indicating the accuracy of in vivo tracer uptake quantification.

### ^99m^Tc-(HE)_3_-(GPO)_9_ autoradiography

Evaluation of in vivo tracer uptake by autoradiography showed significantly higher ^99m^Tc-(HE)_3_-(GPO)_9_ uptake in the infarct zone, detected by Masson’s trichrome and Sirius red staining [0.0026 %ID/cm^2^ (0.0023–0.0035 %ID/cm^2^)] compared to the remote zone of MI-induced mice [0.0016 %ID/cm^2^ (0.0014–0.0024 %ID/cm^2^, n = 6, P < 0.05)], and also compared to the corresponding wall in sham group [(0.0010 %ID/cm^2^ (0.0009–0.0017 %ID/cm^2^), n = 5, P < 0.01] (Fig. 2A and B). Notably, there was a significant correlation between the in vivo, SPECT-based (in %ID/mL) quantification of tracer uptake in the infarct zone and the corresponding anterolateral LV wall in the sham-operated animals and its ex vivo, autoradiography-based (in %ID/mm^2^) quantification of tracer uptake in the same animals, confirming the accuracy of in vivo tracer uptake quantification (Spearman’s R = 0.81, P < 0.01; Fig. 2C).

**Figure 2:**
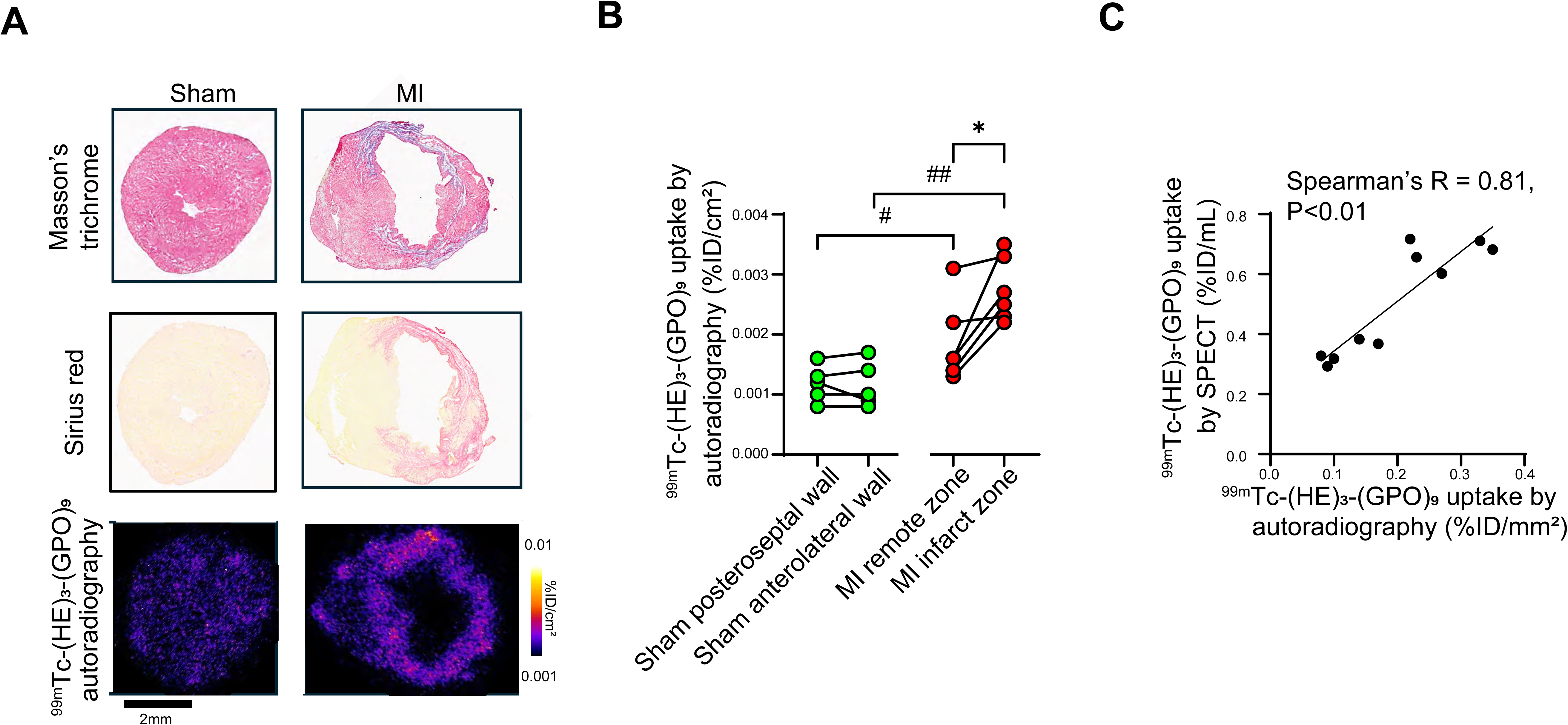
^99m^Tc-(HE)_3_-(GPO)_9_ autoradiography in myocardial infarction. (**A**) Illustrative examples of Masson’s trichrome staining, Sirius red staining, and ^99m^Tc-(HE)_3_-(GPO)_9_ autoradiography (following in vivo tracer administration) of sham-operated and MI-induced mice at two weeks after surgery. (**B**) Quantification of ^99m^Tc-(HE)_3_-(GPO)_9_ uptake by autoradiography in the infarct and remote zones of MI-induced, and the corresponding walls of sham-operated mice. (**C**) Correlation between ^99m^Tc-(HE)_3_-(GPO)_9_ signals on SPECT/CT and autoradiography in the infarct zone and the anterolateral wall of sham-operated mice. ID: injected dose. #: P < 0.05, ##: P < 0.01, Mann-Whitney *U* test; *: P < 0.05, Wilcoxon test.

### Specificity of tracer uptake

To investigate the specificity of ^99m^Tc-(HE)_3_-(GPO)_9_ signal in myocardial infarction, a group of MI-induced animals (n = 7) underwent repeat micro-SPECT/CT-CT imaging with ^99m^Tc-(HE)_3_-(GPO)_9_ and ^99m^Tc-(HE)_3_-(GPO)_SCR_ within 2-3 days. CT imaging with eXIA 160XL, a CT contrast agent that is taken up by viable myocardium (*21*), identified the infarct zone in these animals. The control tracer and ^99m^Tc-(HE)_3_-(GPO)_9_ showed different patterns of uptake throughout the heart (Fig. 3A), and quantification of the infarct zone SPECT signal showed significantly higher uptake of the targeted than the control tracer [0.54 (0.49-0.56 %ID/mL) for ^99m^Tc-(HE)_3_-(GPO)_9_ vs 0.21 (0.20-0.32 %ID/mL) for ^99m^Tc-(HE)_3_-(GPO)_SCR_, P < 0.05, Fig. 3B)], establishing the specificity of the ^99m^Tc-(HE)_3_-(GPO)_9_ signal in myocardial infarction. Evaluation of tracer uptake by autoradiography showed that while there was no difference in in vivo ^99m^Tc-(HE)_3_-(GPO)_SCR_ uptake between the infarct and remote zones of the same hearts, the infarct zone ^99m^Tc-(HE)_3_-(GPO)_SCR_ signal was significantly lower than the ^99m^Tc-(HE)_3_-(GPO)_9_ signal (P < 0.01; Fig. 3C and D).

**Figure 3:**
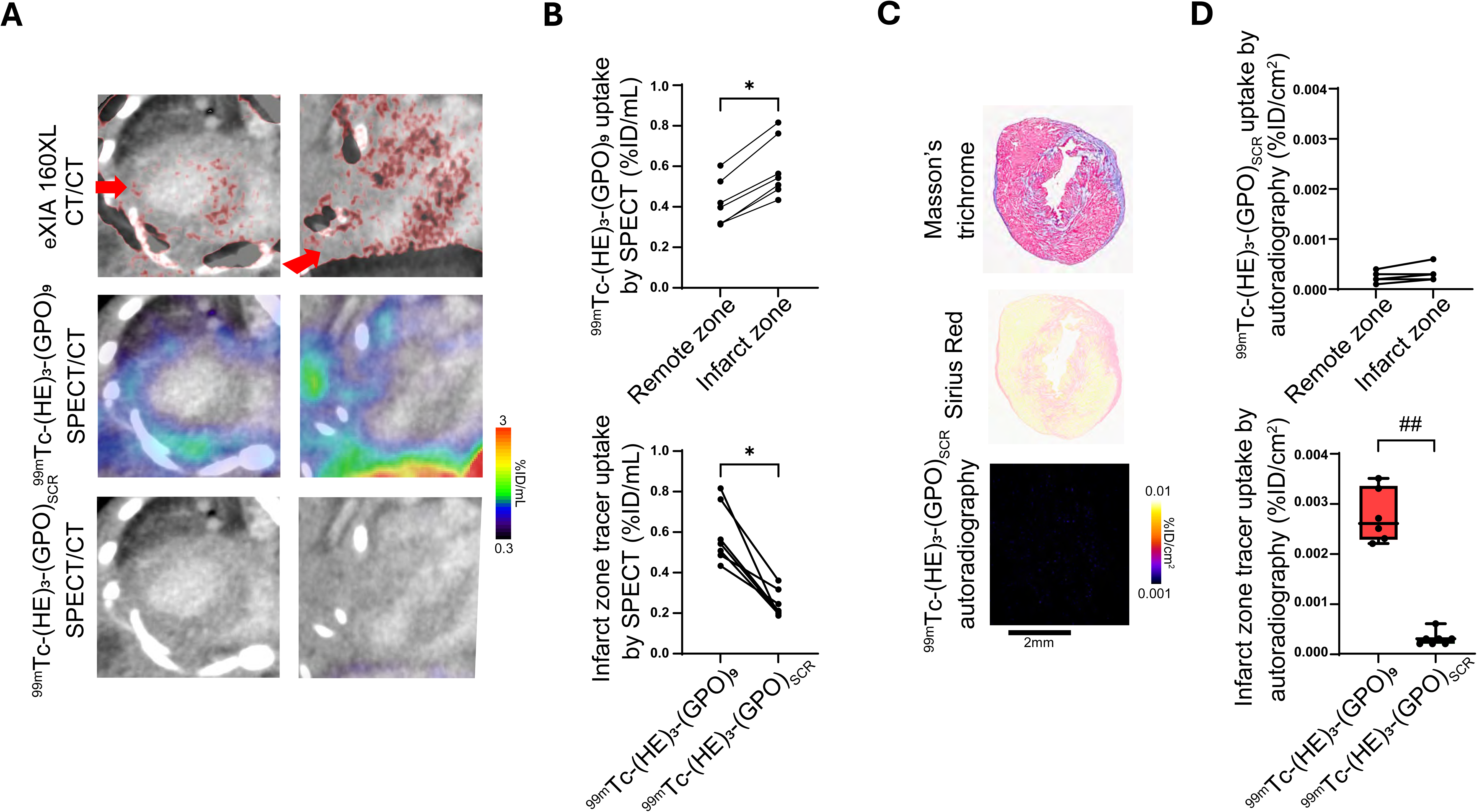
^99m^Tc-(HE)_3_-(GPO)_9_ specificity. (**A**) Illustrative examples of in vivo contrast-enhanced CT using eXIA 160XL fused with an Exitron Nano-12000-enhanced CT (top row), along with SPECT/Exitron Nano-12000-enhanced CT of ^99m^Tc-(HE)_3_-(GPO)_9_ (middle row) and ^99m^Tc-(HE)_3_-(GPO)_SCR_ (bottom row) in MI-induced mice at two weeks after surgery. Red arrows point to the scarred area. (**B**) Quantification of ^99m^Tc-(HE)_3_-(GPO)_9_ SPECT signal in the infarct zone of MI-induced mice in comparison with the ^99m^Tc-(HE)_3_-(GPO)_9_ signal in the remote zone (top panel) and the ^99m^Tc-(HE)_3_-(GPO)_SCR_ signal in the infarct zone (bottom panel) of the same animals. (**C**) Illustrative examples of Masson’s trichrome staining, Sirius red staining, and ^99m^Tc-(HE)_3_-(GPO)_SCR_ autoradiography (following in vivo tracer administration) of MI-induced mice at two weeks after surgery. (**D**) Quantification of ^99m^Tc-(HE)_3_-(GPO)_SCR_ uptake by autoradiography in the infarct zone of MI-induced mice in comparison with ^9m^Tc-(HE)_3_-(GPO)_SCR_ uptake in the remote zone (top panel) and ^99m^Tc-(HE)_3_-(GPO)_9_ uptake in the infarct zone (bottom panel) of the same animals. ID: injected dose. *: P < 0.05, Wilcoxon test; ##: P < 0.01, Mann-Whitney *U* test.

### Correlates of cardiac ^99m^Tc-(HE)_3_-(GPO)_9_ uptake

Evaluation of denatured collagen using a fluorescent collagen hybridizing peptide, R-CHP, in comparison with Sirius red and Masson’s trichrome staining (which detect collagen) showed that denatured collagen is significantly higher in the infarct zone (P < 0.05 when compared to the remote zone or the corresponding wall in sham-operated animals; Fig. 4A and B) at the 2-week timepoint. Notably, there were significant correlations between the denatured collagen quantified by R-CHP staining in the infarct zone and the corresponding anterolateral LV wall in the sham-operated animals, and ^99m^Tc-(HE)_3_-(GPO)_9_ uptake by SPECT/CT (Spearman’s R = 0.78, P < 0.05) and autoradiography (Spearman’s R = 0.82, P < 0.05; Fig. 4C) at 2 weeks post-MI.

**Figure 4:**
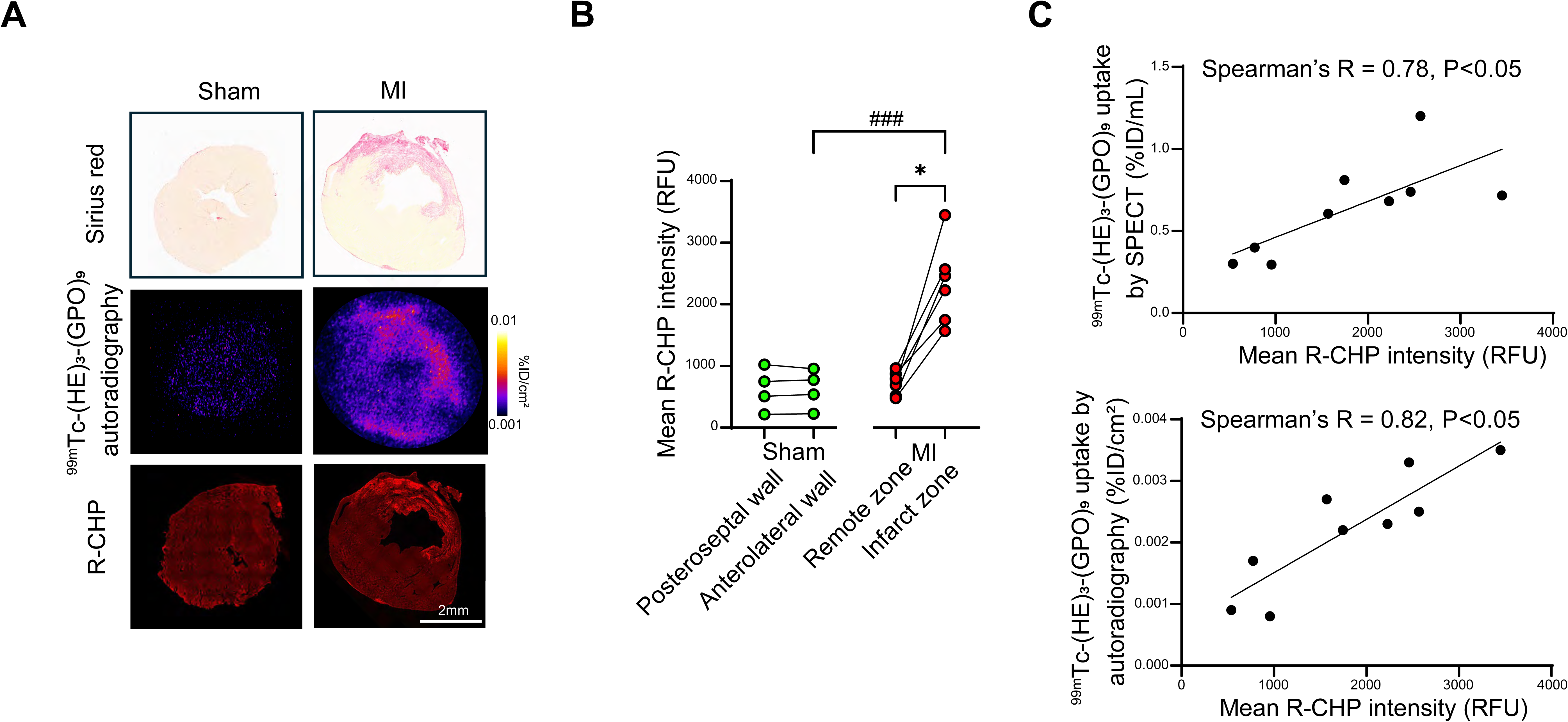
Tracer uptake and denatured collagen. (**A**) Illustrative examples of Sirius red staining, ^99m^Tc-(HE)_3_-(GPO)_9_ autoradiography (following in vivo tracer administration), and denatured collagen staining of sham-operated and MI-induced mice at two weeks after surgery using a fluorescent collagen hybridizing peptide (R-CHP). (**B**) Quantification of R-CHP staining in MI-induced and sham-operated animals. (**C**) Correlation between ^99m^Tc-(HE)_3_-(GPO)_9_ uptake by SPECT/CT (top) or autoradiography (bottom), and R-CHP staining intensity in the infarct zone. ID: injected dose, RFU: relative fluorescence unit. ###: P < 0.001, Mann-Whitney *U* test; *: P < 0.05, Wilcoxon test.

### Denatured collagen and other targets for imaging remodeling

Next, we evaluated the presence of denatured collagen by ex vivo ^99m^Tc-(HE)_3_-(GPO)_9_ autoradiography and R-CHP staining at 3 days, 1 week, and 2 weeks after MI, compared with control animals. Both approaches showed a higher ^99m^Tc-(HE)_3_-(GPO)_9_ signal in the infarct zone, with the ratio of infarct to remote zone increasing over time (Fig. 5). These changes paralleled the changes in MMP activation detected by ex vivo ^99m^Tc-RYM1 autoradiography (*22*). Interestingly, while procollagen expression in the infarct zone was readily detectable at 3 days post-MI, the ratio of denatured collagen (reflecting collagen turnover) to procollagen expression significantly increased from day 3 to 2 weeks post-MI (Supplemental Fig. 5), highlighting the distinct biology of these markers of fibrosis.

**Figure 5:**
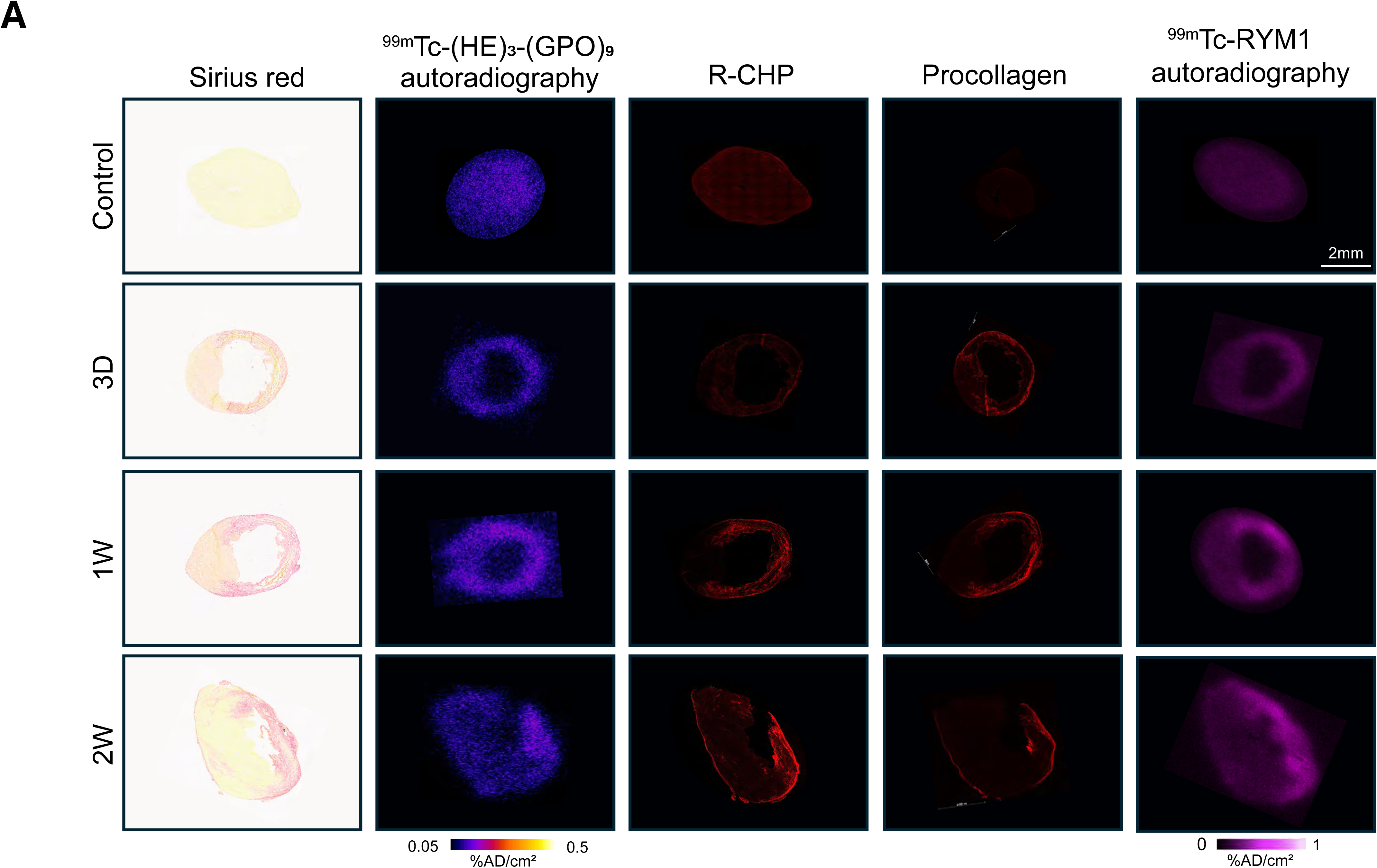
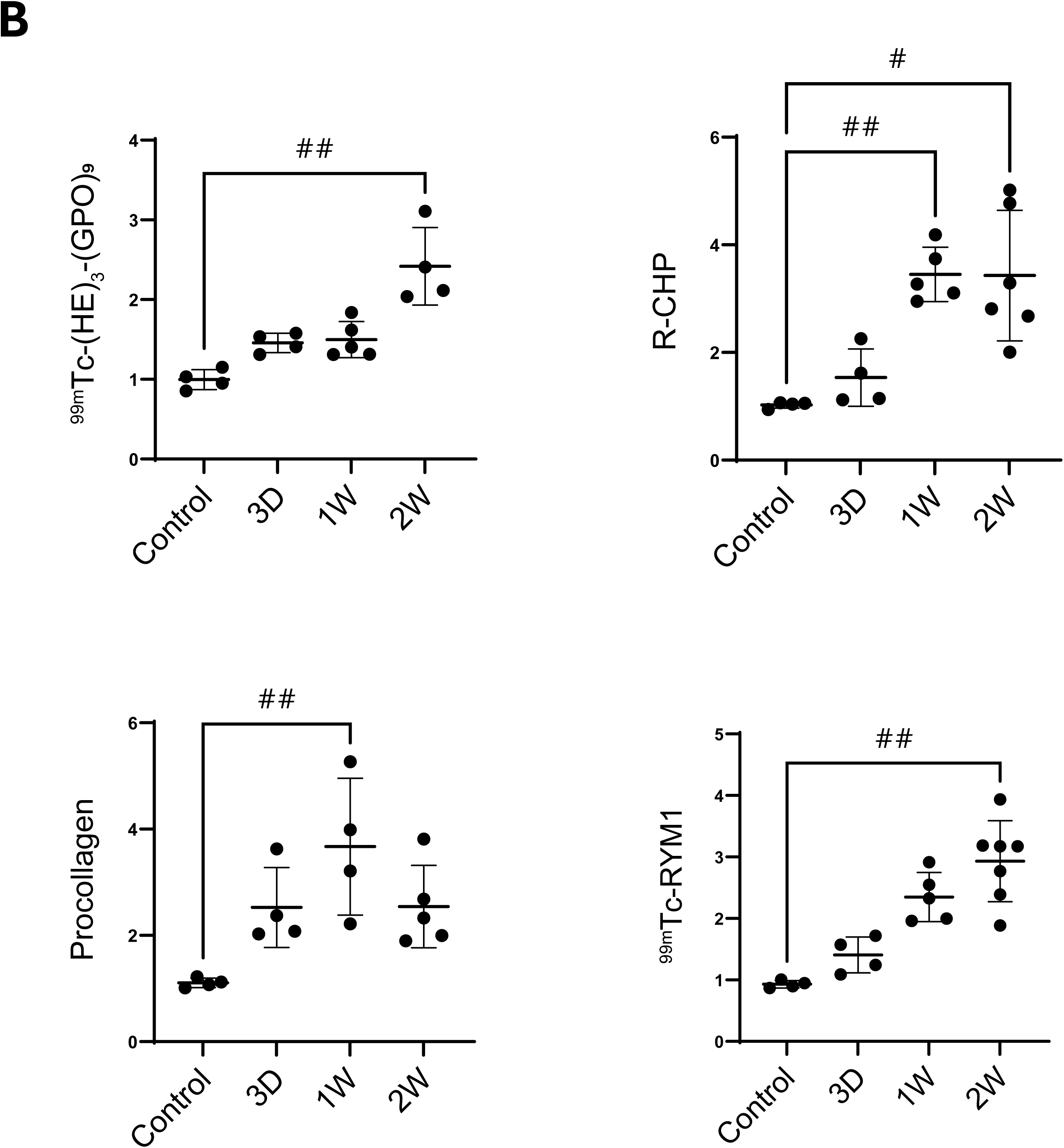
Expression time course of fibrosis-related imaging targets. Illustrative examples (**A**) and infarct-to-remote area signal intensity ratios (**B**) of ex vivo ^99m^Tc-(HE)_3_-(GPO)_9_ autoradiography, R-CHP staining, procollagen staining, and ex vivo ^99m^Tc-RYM1 autoradiography in adjacent sections of the hearts collected from control and 3 days, 1 week, and 2 weeks post-MI animals. Sirius red images are shown as a reference in panel A. AD: applied dose. #: P < 0.05, ##: P < 0.01, Kruskal-Wallis test.

### Human tissue analysis

As a prelude to future human studies, we evaluated the binding of ^99m^Tc-(HE)_3_-(GPO)_9_ to normal and fibrotic human myocardial tissues from patients undergoing myocardial biopsy. The presence of fibrosis was confirmed by Masson’s trichrome and Sirius Red staining. The normal myocardium showed little denatured collagen as detected by R-CHP staining. In contrast, R-CHP staining of the fibrotic biopsies showed a heterogeneous pattern, with denatured collagen localized to fibrotic regions identified by Masson’s trichrome and Sirius red staining. Similarly, while there was minimal ^99m^Tc-(HE)_3_-(GPO)_9_ binding to the normal myocardium, the fibrotic biopsies showed a heterogeneous pattern of ^99m^Tc-(HE)_3_-(GPO)_9_ binding localized to the fibrotic areas (Fig. 6).

**Figure 6:**
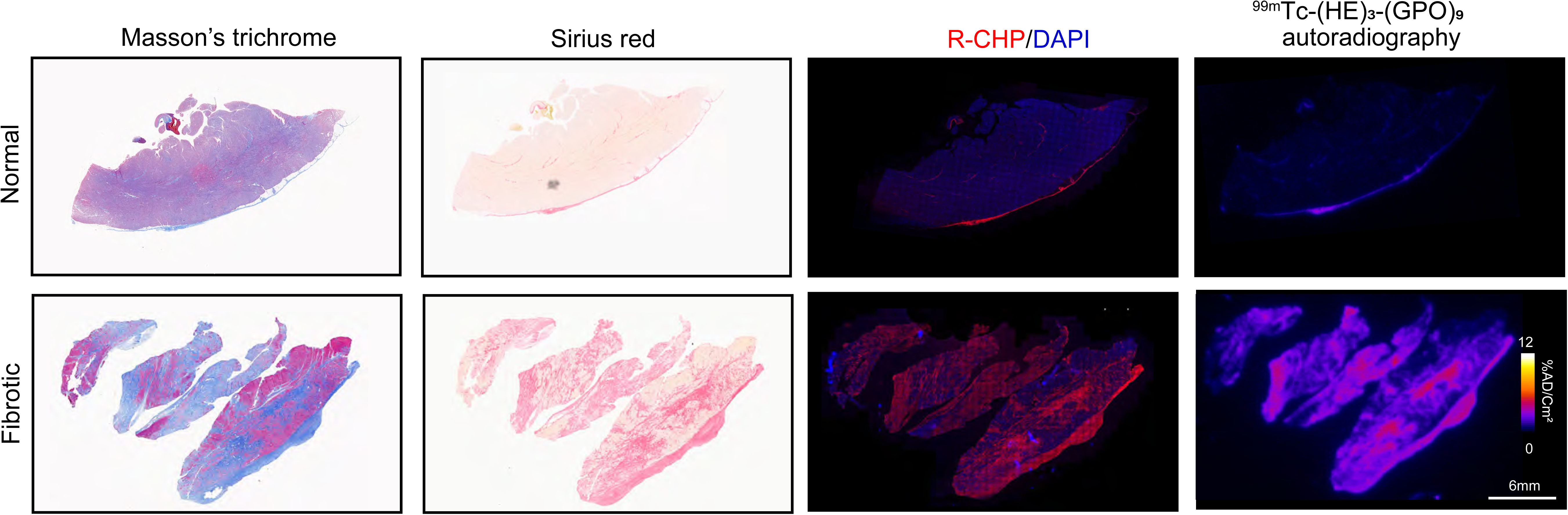
^99m^Tc-(HE)_3_-(GPO)_9_ binding to human myocardial tissue. Illustrative examples of Masson’s trichrome staining of fibrosis, Sirius red staining of collagen, denatured collagen staining using a fluorescent collagen hybridizing peptide (R-CHP), and ^99m^Tc-(HE)_3_-(GPO)_9_ autoradiography in normal and fibrotic myocardium.

## Discussion

Using a recently introduced collagen hybridizing tracer, ^99m^Tc-(HE)_3_-(GPO)_9_, we demonstrated the feasibility of in vivo imaging of denatured collagen as a marker of collagen turnover following MI. SPECT/CT images showed higher tracer uptake in the infarct zone compared to the remote zone in animals with MI, and in the corresponding LV region in sham-operated animals. The accuracy of in vivo signal quantification was established by comparison with ex vivo autoradiography and gamma-counting, and the specificity of tracer uptake was confirmed by comparison with the uptake of a control, scrambled homolog tracer. Evaluation of denatured collagen by histology showed a strong correlation with in vivo tracer uptake. Notably, various tissue markers of fibrosis exhibited distinct spatial patterns in MI, suggesting their complementarity as molecular imaging targets. Finally, we demonstrated that the tracer binds effectively to denatured human collagen in fibrotic myocardial biopsies.

Fibrosis is a key characteristic of maladaptive ventricular remodeling, i.e., cardiomyopathy, that underlies the development of most heart failure cases. It is often triggered by cardiac injury and mediated by activated cardiac fibroblasts that deposit fibrotic tissue within the myocardium. The response to injury involves inflammation, fibroblast proliferation, and the transformation of these cells into myofibroblasts, ultimately leading to fibrosis and scar formation (*23*). Classically, cardiac fibrosis is classified into reactive interstitial fibrosis and replacement fibrosis (*23*). Whether interstitial or replacement, cardiac fibrosis directly contributes to structural and biological changes that eventually result in heart failure.

Replacement fibrosis is the initial response to MI, where there is extensive cardiomyocyte death, and the repair process seeks to prevent cardiac rupture. This repair process follows the immediate phase of inflammatory cell recruitment and eventually may be associated with remote-zone reactive interstitial fibrosis and maladaptive ventricular remodeling (*24*). By altering the mechano-electric coupling of cardiomyocytes, cardiac fibrosis also promotes arrhythmogenesis (*23*) and, through paracrine effects, influences myocyte biology (*25*). As such, cardiac fibrosis directly contributes to heart failure and its complications. Given the role of fibrosis in cardiac remodeling and arrhythmias, there is considerable interest in new therapies that can prevent or reverse cardiac fibrosis (*23*). The success of these efforts is dependent on access to non-invasive, quantitative tools to track the development and resolution of cardiac fibrosis and ventricular remodeling.

Typically, the diagnosis of cardiac fibrosis is based on (gadolinium-enhanced) magnetic resonance imaging (MRI). Alternative approaches include myocardial perfusion imaging and other nuclear imaging techniques to detect scar, and integrated backscatter by ultrasound (*26*). However, these techniques provide a snapshot of the cardiac structure at a given point, without providing any information on disease *activity*, which is arguably the primary target of therapeutic interventions aimed at preventing progression and promoting regression of fibrosis. Additionally, there is a lag before the effects of therapeutic interventions are reflected in cardiac structure and function, and thus become detectable by MRI or ultrasound. Accordingly, novel tools are needed to detect collagen turnover during fibrosis, select patients for emerging therapies, track the effects of therapeutic interventions to guide their timing, and potentially improve prognostication.

To address this unmet need, several new tracers for molecular imaging of fibrosis-related processes have been introduced in recent years. These agents target integrins, fibroblasts, or matrix components such as mature collagen (*27*). However, none of these techniques can distinguish between established disease and ongoing matrix remodeling, which accompanies active fibrogenesis and resolution of fibrosis, i.e., collagen remodeling (turnover) (*27*). Gadolinium-labeled EP-3533 and CM-101 for MRI, ^68^Ga-CBP8 for PET imaging, and ^99m^Tc-collagelin and ^99m^Tc-CBP1495 for SPECT imaging are examples of probes under development that bind to mature collagen fibers (*27*). Other emerging tracers target collagen cross-linking, a key step in collagen fiber maturation (*28,29*). However, neither of these approaches provides adequate information on collagen denaturation and degradation, a key component of ECM remodeling, and a target of therapeutic interventions in fibrosis. Emerging techniques for imaging the activation of proteases involved in tissue remodeling, such as MMPs (*20,22,30,31*), may in part address these issues. However, at best, they would provide an *indirect* measure of the downstream effect, i.e., matrix remodeling.

The hallmark of collagen structure is the triple helix, a right-handed helix of 3 α-chains. α-chains are formed by repetitive glycine (Gly)-X-Y (where X and Y are frequently proline and hydroxyproline) tri-peptide motifs, which self-assemble to form (pro)collagen fibers (*32*). Collagen is initially synthesized as procollagen, a precursor molecule with C-and N-terminal propeptides flanking the Gly-X-Y motifs. Following translation, the procollagen α-chains assemble into triple helix procollagen. To form collagen, procollagen undergoes proteolytic cleavage through the removal of the N- and C-terminal propeptides following its export into the extracellular space, where it assembles into fibrils and higher-ordered structures (*32*). The cross-linking of lysine and hydroxylysine residues within and between collagen molecules, facilitated by lysyl oxidase, strengthens the collagen fibrils. The α-chains are highly organized in mature collagen fibers, and single or double-stranded α-chains are only present in the extracellular space when collagen is undergoing degradation, e.g., by matrix metalloproteinases (MMPs) and cathepsins (*32*). Collagen hybridizing peptides bind to single- or double-stranded collagen and may be used to detect the degraded and unfolded collagen chains through triple helix formation (*16*).

Little information is available regarding MI-induced spatial and temporal changes in denatured collagen. Our evaluation of denatured collagen post-MI using a fluorescent CHP showed an early increase in denatured collagen in the infarct zone compared to the remote zone, which augmented from 3 days to 2 weeks. Later changes in denatured collagen in both the infarct and remote zones, and their relation to ventricular remodeling, could be a subject of future studies. Notably, there were considerable differences in denatured collagen content between different animals, especially at later time points, which potentially reflects the variability of the response to LAD ligation in this murine model, and highlights the value of longitudinal studies to establish the relationship between denatured collagen and LV structure and function, in the absence or presence of standard post-MI and emerging anti-fibrotic therapies.

As a key step toward such studies, we demonstrated the feasibility of in vivo imaging of denatured collagen as a marker of post-MI collagen turnover. SPECT/CT images showed higher tracer uptake in the infarct zone compared to the remote zone in animals with MI, and in the corresponding LV region in sham-operated animals. The specificity of tracer uptake was confirmed by comparing it with the uptake of a control, scrambled homolog tracer. The tracer uptake in the surgical wound and the liver made quantification of cardiac tracer uptake somewhat challenging. Based on images obtained with the control tracer, the targeted tracer uptake in the wound was specific, reflecting collagen remodeling during wound healing. Similarly, a large component of liver uptake appeared to be specific, potentially reflecting growth-related remodeling in these young animals. Despite these challenges, the accuracy of in vivo signal quantification was established through comparison with ex vivo autoradiography and gamma-counting. Furthermore, histological evaluation of denatured collagen showed a strong correlation with in vivo tracer uptake. Together, these studies established the feasibility and validity of ^99m^Tc-(HE)_3_-(GPO)_9_ SPECT/CT for detecting denatured collagen post-MI. The binding of ^99m^Tc-(HE)_3_-(GPO)_9_ to human denatured collagen in fibrotic myocardial biopsies, although not unexpected given the similarity of collagen structure across species, supported the potential of this family of tracers for imaging in humans.

The similarity between the spatial and temporal patterns of denatured collagen and MMP activation, as revealed by RYM1 autoradiography, highlights the role of MMPs in collagen remodeling. Based on these results, one can speculate that imaging MMP and GPO imaging will provide similar information regarding post-MI ventricular remodeling. Whether this similarity persists in an antifibrotic therapy setting remains to be determined. The distinct patterns of procollagen expression and denatured collagen, on the other hand, reflect their distinct biology in the development and resolution of fibrosis. The significant increase in the ratio of denatured collagen to procollagen expression from 3 days to 2 weeks post-MI indicates a shift from collagen production to collagen remodeling within this period. Fibroblast activation, an early step in collagen production, can be imaged using emerging FAPI tracers. Our data suggests that combining imaging of denatured collagen with FAP imaging may provide complementary information regarding post-MI fibrosis.

There are several limitations to this initial report, which focuses on establishing the potential of CHP-based tracers for imaging denatured collagen after MI. As such, the long-term changes in collagen denaturation, the functional significance of the denatured collagen signal post-MI, and the effect of anti-fibrotic therapies on the CHP signal remain to be addressed in future studies. Although validation in a mouse model facilitated future mechanistic studies of collagen denaturation using genetically modified mice, the small size of the mouse LV limited our ability to evaluate regional changes in imaging targets in more detail. In this regard, the detection and quantification of the signal in the heart would be facilitated by developing tracers with reduced non-specific liver uptake.

## Conclusion

^99m^Tc-(HE)_3_-(GPO)_9_ imaging of denatured collagen adds a new dimension to fibrosis imaging by complementing other emerging approaches that target the development and resolution of fibrosis and ventricular remodeling. Access to this approach enables more in-depth mechanistic studies on collagen turnover in cardiomyopathy and can help develop and validate new antifibrotic therapies. The clinical translation of our findings should ultimately lead to improved management of cardiomyopathy.

## KEY POINTS

QUESTION: Can we image denatured collagen as a marker of collagen remodeling post-myocardial infarction?

PERTINENT FINDINGS: SPECT/CT using ^99m^Tc-(HE)_3_-(GPO)_9_, a collagen-hybridizing radiotracer, showed higher specific uptake in the infarct zone of MI mice than in the remote zone and sham controls. Evaluation of collagen production and denaturation after myocardial infarction revealed a transition from collagen synthesis to remodeling from 3 days to 2 weeks post-infarction.

IMPLICATIONS FOR PATIENT CARE: ^99m^Tc-(HE)_3_-(GPO)_9_ enables the detection of collagen remodeling after myocardial infarction, filling a gap in biological and clinical research that could ultimately improve post-infarction risk stratification and management by monitoring the effect of emerging anti-fibrotic therapies.

---

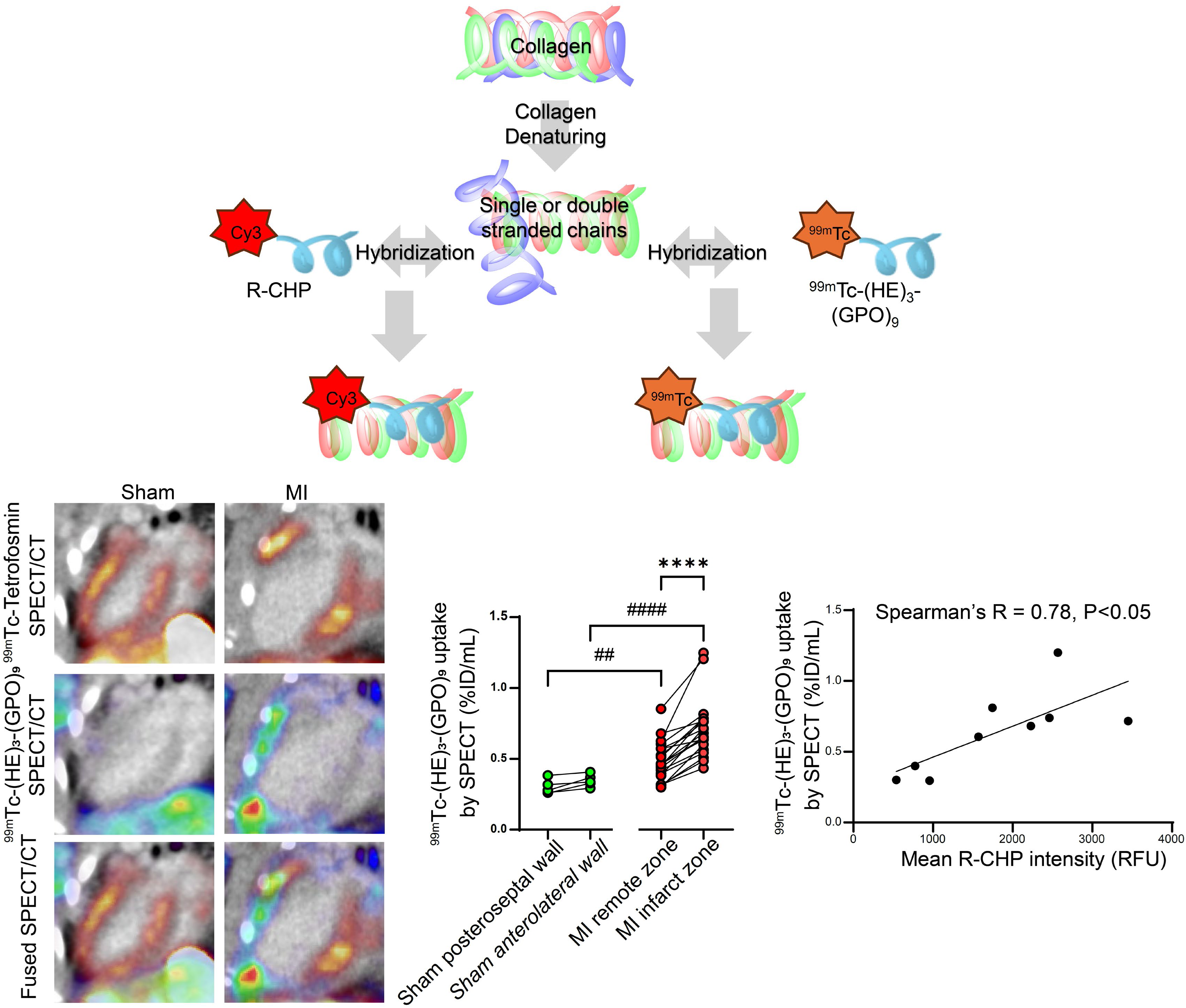

## References

1. Wynn TA, Ramalingam TR. Mechanisms of fibrosis: therapeutic translation for fibrotic disease. Nat Med. 2012;18:1028–1040.

2. Henderson NC, Rieder F, Wynn TA. Fibrosis: from mechanisms to medicines. Nature. 2020;587:555–566.

3. Ezeani M, Noor A, Alt K, et al. Collagen-Targeted Peptides for Molecular Imaging of Diffuse Cardiac Fibrosis. J Am Heart Assoc. 2021;10:e022139.

4. Frangogiannis NG. Cardiac fibrosis. Cardiovasc Res. 2021;117:1450–1488.

5. Bahit MC, Kochar A, Granger CB. Post-Myocardial Infarction Heart Failure. JACC Heart Fail. 2018;6:179–186.

6. Frantz S, Hundertmark MJ, Schulz-Menger J, Bengel FM, Bauersachs J. Left ventricular remodelling post-myocardial infarction: pathophysiology, imaging, and novel therapies. Eur Heart J. 2022;43:2549–2561.

7. Mangion K, McComb C, Auger DA, Epstein FH, Berry C. Magnetic Resonance Imaging of Myocardial Strain After Acute ST-Segment-Elevation Myocardial Infarction: A Systematic Review. Circ Cardiovasc Imaging. 2017;10.

8. Arun S, Mittal BR, Bhattacharya A, Rohit MK. Comparison of Tc-99m tetrofosmin myocardial perfusion scintigraphy and exercise F18-FDG imaging in detection of myocardial ischemia in patients with coronary artery disease. J Nucl Cardiol. 2015;22:98–110.

9. Makowski MR, Rischpler C, Ebersberger U, et al. Multiparametric PET and MRI of myocardial damage after myocardial infarction: correlation of integrin alphavbeta3 expression and myocardial blood flow. Eur J Nucl Med Mol Imaging. 2021;48:1070–1080.

10. Muzard J, Sarda-Mantel L, Loyau S, et al. Non-invasive molecular imaging of fibrosis using a collagen-targeted peptidomimetic of the platelet collagen receptor glycoprotein VI. PLoS One. 2009;4:e5585.

11. Nahrendorf M, Hu K, Frantz S, et al. Factor XIII deficiency causes cardiac rupture, impairs wound healing, and aggravates cardiac remodeling in mice with myocardial infarction. Circulation. 2006;113:1196–1202.

12. Niego B, Jupp B, Zia NA, et al. Molecular Imaging of Diffuse Cardiac Fibrosis with a Radiotracer That Targets Proteolyzed Collagen IV. Radiol Cardiothorac Imaging. 2024;6:e230098.

13. Sahul ZH, Mukherjee R, Song J, et al. Targeted imaging of the spatial and temporal variation of matrix metalloproteinase activity in a porcine model of postinfarct remodeling: relationship to myocardial dysfunction. Circ Cardiovasc Imaging. 2011;4:381–391.

14. Varasteh Z, Mohanta S, Robu S, et al. Molecular Imaging of Fibroblast Activity After Myocardial Infarction Using a (68)Ga-Labeled Fibroblast Activation Protein Inhibitor, FAPI-04. J Nucl Med. 2019;60:1743–1749.

15. Meoli DF, Sadeghi MM, Krassilnikova S, et al. Noninvasive imaging of myocardial angiogenesis following experimental myocardial infarction. J Clin Invest. 2004;113:1684–1691.

16. Bennink LL, Li Y, Kim B, et al. Visualizing collagen proteolysis by peptide hybridization: From 3D cell culture to in vivo imaging. Biomaterials. 2018;183:67–76.

17. Li X, Zhang Q, Yu SM, Li Y. The Chemistry and Biology of Collagen Hybridization. J Am Chem Soc. 2023;145:10901–10916.

18. Li Y, Foss CA, Summerfield DD, et al. Targeting collagen strands by photo-triggered triple-helix hybridization. Proc Natl Acad Sci U S A. 2012;109:14767–14772.

19. Ahmad AA, Ghim M, Kukreja G, et al. Collagen Hybridizing Peptide-Based Radiotracers for Molecular Imaging of Collagen Turnover in Pulmonary Fibrosis. J Nucl Med. 2025;66:425–433.

20. Su H, Spinale FG, Dobrucki LW, et al. Noninvasive targeted imaging of matrix metalloproteinase activation in a murine model of postinfarction remodeling. Circulation. 2005;112:3157–3167.

21. Ashton JR, Befera N, Clark D, et al. Anatomical and functional imaging of myocardial infarction in mice using micro-CT and eXIA 160 contrast agent. Contrast Media Mol Imaging. 2014;9:161–168.

22. Toczek J, Ye Y, Gona K, et al. Preclinical Evaluation of RYM1, a Matrix Metalloproteinase-Targeted Tracer for Imaging Aneurysm. J Nucl Med. 2017;58:1318–1323.

23. Travers JG, Kamal FA, Robbins J, Yutzey KE, Blaxall BC. Cardiac Fibrosis: The Fibroblast Awakens. Circ Res. 2016;118:1021–1040.

24. Beltrami CA, Finato N, Rocco M, et al. Structural basis of end-stage failure in ischemic cardiomyopathy in humans. Circulation. 1994;89:151–163.

25. Kakkar R, Lee RT. Intramyocardial fibroblast myocyte communication. Circ Res. 2010;106:47–57.

26. Jellis C, Martin J, Narula J, Marwick TH. Assessment of nonischemic myocardial fibrosis. J Am Coll Cardiol. 2010;56:89–97.

27. Montesi SB, Desogere P, Fuchs BC, Caravan P. Molecular imaging of fibrosis: recent advances and future directions. J Clin Invest. 2019;129:24–33.

28. Shuvaev S, Knipe RS, Drummond M, et al. Optimization of an Allysine-Targeted PET Probe for Quantifying Fibrogenesis in a Mouse Model of Pulmonary Fibrosis. Mol Imaging Biol. 2023;25:944–953.

29. Wahsner J, Desogere P, Abston E, et al. (68)Ga-NODAGA-Indole: An Allysine-Reactive Positron Emission Tomography Probe for Molecular Imaging of Pulmonary Fibrogenesis. J Am Chem Soc. 2019;141:5593–5596.

30. Zhang J, Nie L, Razavian M, et al. Molecular imaging of activated matrix metalloproteinases in vascular remodeling. Circulation. 2008;118:1953–1960.

31. Gona K, Toczek J, Ye Y, et al. Hydroxamate-Based Selective Macrophage Elastase (MMP-12) Inhibitors and Radiotracers for Molecular Imaging. J Med Chem. 2020;63:15037–15049.

32. Mouw JK, Ou G, Weaver VM. Extracellular matrix assembly: a multiscale deconstruction. Nat Rev Mol Cell Biol. 2014;15:771–785.

