## Supplemental Data for "Molecular Imaging of Collagen Turnover in Myocardial Infarction"

[*] Equal contribution

### Supplemental Material

#### Table of Contents


| | |
|---|---|
| Detailed Materials and Methods | 3 |
| Supplemental Figure 1: Flowchart of experiments | 7 |
| Supplemental Figure 2: Tracer quality control | 8 |
| Supplemental Figure 3: Impact of surgical procedures on cardiac function | 9 |
| Supplemental Figure 4: Biodistribution of $^{99m}Tc$-$(HE)_3$-$(GPO)_9$ | 10 |
| Supplemental Figure 5: Collagen production and denaturation | 11 |

#### Supplemental Materials and Methods

##### Radiotracers

The $(HE)_3$-$(GPO)_9$ precursor was synthesized and radiolabeled to generate $^{99m}Tc$-$(HE)_3$-$(GPO)_9$ as described (*1*). This tracer, which targets denatured collagen, features a polyhistidine–glutamic acid [$(HE)_3$] N-terminal sequence for site-specific radiolabeling, linked to a C-terminal targeting moiety consisting of 9 GPO repeats [$(GPO)_9$] via a flexible 3-glycine linker. A control tracer, $^{99m}Tc$-$(HE)_3$-$(GPO)_{SCR}$, where $(GPO)_9$ is replaced with a scrambled sequence (PGOGPGPOPOGOGOPPGOOPGGOOPPG), was used as a negative control agent. RYM1, a macrocyclic hydroxamate-based pan-MMP tracer precursor, was labeled with $^{99m}Tc$ to give $^{99m}Tc$-RYM1 (specific activity of 264±33 GBq/μmol) as previously described (*2*). The identity and purity of precursors were confirmed by liquid chromatography–mass spectrometry (LC-MS). The purity of tracers was confirmed by radio-high-performance liquid chromatography (Radio-HPLC). Tetrofosmin (Mediphysics Amersham Healthcare, Arlington Heights, IL) was labeled with $^{99m}Tc$ according to standard protocols (*3*).

##### Animals

Animal experiments were performed under protocols approved by the Institutional Animal Use and Care Committees of Yale University and the Veterans Affairs Connecticut Healthcare System. To induce MI, 10- to 16-week-old C57BL/6J mice (n = 39, Supplemental Fig. 1) of both sexes underwent left anterior descending (LAD) artery ligation 2 mm distal to the left auricle, as described with some modifications (*4*). Briefly, under isoflurane anesthesia (1-3% in oxygen), the animals were intubated in the supine position and ventilated at a tidal volume of 0.2 mL and a rate of 120 cycles/min. Using a sterile technique, a left thoracotomy in the fourth intercostal space was performed, and the pericardium was opened. The left anterior descending coronary artery was ligated with a 7-0 monofilament suture, and the incision was closed in two layers. The surgical mortality rate (either because of sudden death or a requirement for euthanasia because of deteriorating status) over 2 weeks was 28%. Sham-operated or unoperated animals (n = 8, Supplemental Fig. 1) were used as controls.

##### Human tissues

All experiments were conducted in accordance with protocols approved by Yale University. Fibrotic and non-involved heart tissues were from anonymized patients who had undergone myocardial biopsy. Tissues were provided as formalin-fixed, paraffin-embedded tissue sections. Adjacent sections were used for autoradiography, Sirius red staining, and Masson's trichrome staining.

**Echocardiography**

Echocardiographic images were obtained as reported previously (*5*). Briefly, echocardiography was performed using a VisualSonics Vevo 2100 system with a 40 MHz MS550D transducer. Anesthesia was induced with 1.5–2.5% isoflurane. A rectal probe was used to monitor the body temperature. Long axis B mode measurements were performed to assess left ventricular ejection fraction (LVEF). One animal was excluded due to an unrealistically low calculated EF.

**SPECT/CT imaging**

A subset of animals underwent $^{99m}$Tc-tetrofosmin myocardial perfusion imaging (MPI) 2 weeks after MI or a sham operation (Supplemental Fig. 1). Animals were anesthetized under 1–3% isoflurane, retro-orbitally injected with 26.3±4.1 MBq of $^{99m}$Tc-tetrofosmin and imaged after 20 minutes. SPECT image acquisition was performed on a small-animal SPECT/CT scanner (U-SPECT4CT, MILabs) for 15 minutes and was followed by a full-body CT scan, as described (*1*). Animals were allowed to rest for 2-3 days following tetrofosmin imaging to allow the radiotracer to clear before $^{99m}$Tc-(HE)$_3$-(GPO)$_9$ imaging. $^{99m}$Tc-(HE)$_3$-(GPO)$_9$ SPECT/CT imaging was performed 1 hour after retro-orbital injection of 29.2±7.8 MBq of heated (to 80 °C for 10 minutes) and cooled (on ice for 30 seconds) $^{99m}$Tc-(HE)$_3$-(GPO)$_9$ and 30 µL + body weight in grams µL of Exitron Nano-12000 (Viscover). List-mode images were obtained for 45 minutes. Instead of tetrofosmin MPI, another subset of animals was injected retro-orbitally with 50 µL + body weight in grams µL of eXIA 160XL (Binitio Biomedical Inc.), a CT contrast agent that is taken up by viable myocardium (*6*), and imaged by CT 210-270 minutes later to identify the infarct zone. To address tracer uptake specificity, a subset of animals underwent $^{99m}$Tc-(HE)$_3$-(GPO)$_9$ SPECT (23.3±6.7 MBq) imaging and $^{99m}$Tc-(HE)$_3$-(GPO)$_{SCR}$ SPECT (26.3±6.3 MBq) within 2-3 days. Each SPECT study was coupled with either eXIA 160XL- or Exitron Nano-12000-enhanced CT.

The MILabs software (version 12) was used to reconstruct contrast-enhanced (CE)-CT images at an isotropic voxel size of 0.1 mm using a filtered back-projection algorithm. Emission data were reconstructed using a similarity-regulated ordered-subsets expectation maximization algorithm with a 0.4-mm voxel size, 16 subsets, 9 iterations, and a 0.45-mm full-width-at-half-maximum Gaussian filter. The SPECT data were then registered to the CE-CT images, and the images were quantified using 3D Slicer software (version 5.2.2; https://www.slicer.org/) and visualized with AMIDE software (version 1.0.6;

https://amide.sourceforge.net/). Segmentations of infarct and remote zones, or their corresponding LV walls, were performed based on tetrofosmin MPI or eXIA 160XL CE-CT images and applied to SPECT images to measure tracer uptake as the average %injected dose (ID)/mL.

**Biodistribution and autoradiography**

Following SPECT/CT imaging and at two hours after tracer injection, a blood sample was obtained, and the animals were euthanized. Tissue samples were collected from the heart apex, and several other organs, and their radioactivity was quantified using a gamma-well counter (Wizard2; PerkinElmer), as percentage %ID/g along with blood's activity as %ID/mL. The remaining heart was then placed in optimal cutting temperature (OCT), stored on dry ice, and sectioned on the same day. For quantitative autoradiography, the heart was sectioned at 10 μm thickness in the apical and basal regions. The sections and references of known activity were exposed on a phosphor screen (MultiSensitive Phosphor Screen; PerkinElmer) and subsequently scanned with a phosphor imager (Typhoon Trio; GE Healthcare Life Sciences). The injected dose was calibrated to a set time to account for decay from the time of injection to the time of exposure. The tracer signals in the remote and infarct zones (and corresponding walls of sham-operated animals) were measured on calibrated images using Fiji/ImageJ software (National Institutes of Health), and the results were expressed as percentage injected dose (%ID) per square centimeter.

For human tissue autoradiography, the slides were deparaffinized and rehydrated in xylene for 10 minutes twice, then 100% ethanol for 10 minutes twice, followed by 75% ethanol for 5 minutes and 50% ethanol for 5 minutes. Finally, they were rinsed in PBS. For mouse heart autoradiography, 7 μm-thick tissue sections were fixed in 10% neutral buffered formalin (NBF). All sections were blocked in 5% bovine serum albumin, incubated with 0.57±0.04 MBq (in 600-800 μL) of $^{99m}$Tc-(HE)$_3$-(GPO)$_9$ at 37 °C or $^{99m}$Tc-RYM1 at RT for 60 min, and then washed in PBS for 5 min x 3. The slides and known radioactivity standards were exposed to a phosphor plate overnight and scanned with a phosphor imager. Trichrome images were analyzed in ImageJ to generate a binary mask of blue collagen, which was then overlaid onto autoradiography.

**Histology and immunostaining**

Yale Pathology Tissue Services performed Sirius red and Masson's trichrome staining on 7 μm-thick, 10% NBF-fixed sections according to standard protocols. Staining with a fluorescent sulfo-cyanine-labeled CHP (R-CHP, 10 μM; Product No: RED300 / RED60; 3Helix) and procollagen

(ThermoFisher, #PA5-35380, with Alexa Fluor 594 donkey anti-rabbit IgG, Abcam, #A-21207 as secondary) was performed on adjacent tissue sections and analyzed as described (*1*). Regions of interest were drawn over the infarct and remote zones based on Sirius red staining for each tissue section (or corresponding walls in sham-operated animals), and the mean intensity was recorded.

**Statistical analysis**

Descriptive data are reported as median and interquartile range (Q1–Q3) or mean and SD, as appropriate. Paired groups were compared using the Wilcoxon matched-pairs test. Unpaired groups were compared using the Mann-Whitney *U* test. Multiple groups were compared using the Kruskal-Wallis test. The correlation between two variables was determined using Spearman's rank-order correlation. Outlier data points, determined using GraphPad's Robust Regression and Outlier Removal (ROUT, Q = 1%) method, were removed from the SPECT and correlation analyses. Statistical analyses were performed using GraphPad Prism version 10 (GraphPad Software) or later, and a P value of less than 0.05 was considered significant.

#### Supplemental References

**1.** Ahmad AA, Ghim M, Kukreja G, et al. Collagen Hybridizing Peptide-Based Radiotracers for Molecular Imaging of Collagen Turnover in Pulmonary Fibrosis. *J Nucl Med.* 2025;66:425-433.

**2.** Toczek J, Ye Y, Gona K, et al. Preclinical Evaluation of RYM1, a Matrix Metalloproteinase-Targeted Tracer for Imaging Aneurysm. *J Nucl Med.* 2017;58:1318-1323.

**3.** Jain D, Wackers FJ, Mattera J, McMahon M, Sinusas AJ, Zaret BL. Biokinetics of technetium-99m-tetrofosmin: myocardial perfusion imaging agent: implications for a one-day imaging protocol. *J Nucl Med.* 1993;34:1254-1259.

**4.** Su H, Spinale FG, Dobrucki LW, et al. Noninvasive targeted imaging of matrix metalloproteinase activation in a murine model of postinfarction remodeling. *Circulation.* 2005;112:3157-3167.

**5.** Ahmad AA, Ghim M, Toczek J, et al. Multimodality Imaging of Aortic Valve Calcification and Function in a Murine Model of Calcific Aortic Valve Disease and Bicuspid Aortic Valve. *J Nucl Med.* 2023;64:1487-1494.

**6.** Ashton JR, Befera N, Clark D, et al. Anatomical and functional imaging of myocardial infarction in mice using micro-CT and eXIA 160 contrast agent. *Contrast Media Mol Imaging.* 2014;9:161-168.

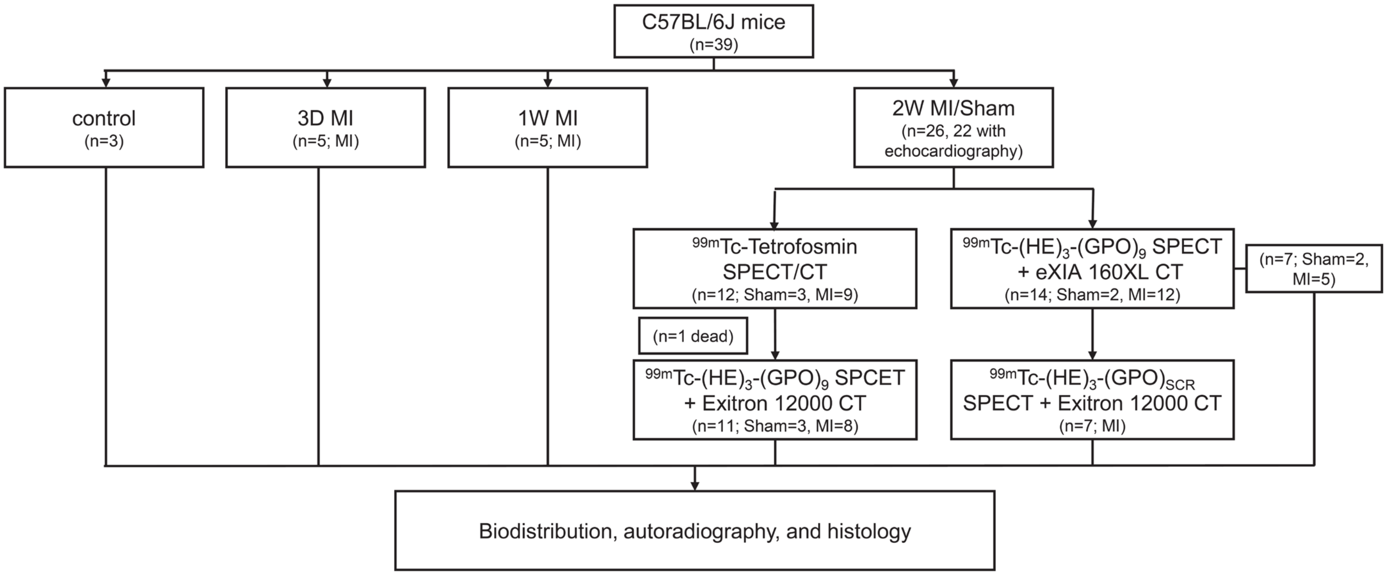


**Supplemental Figure 1: Flowchart of experiments.** Schematic description of experimental procedures performed with the number of mice used for each study. In animals undergoing SPECT imaging with both targeted and control tracers, eXIA 160XL CT and Exitron 12000 CT were performed in different orders. MI: Myocardial infarction.

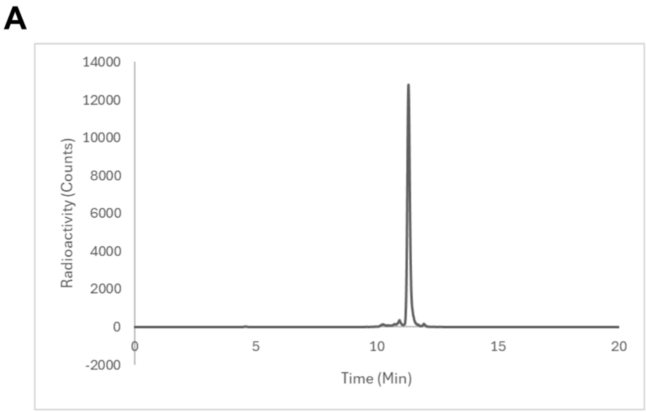


**Supplemental Figure 2: Tracer quality control.** Representative radio-HPLC profiles of $^{99m}$Tc-(HE)$_3$-(GPO)$_9$ (**A**) and $^{99m}$Tc-(HE)$_3$-(GPO)$_{SCR}$ (**B**).

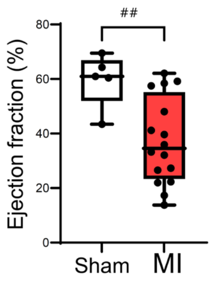


**Supplemental Figure 3: Impact of surgical procedures on cardiac function.** Left ventricular ejection fraction measured by echocardiography in MI-induced and sham-operated animals at 2 weeks post-surgery. ##: P < 0.01, Mann-Whitney *U* test.

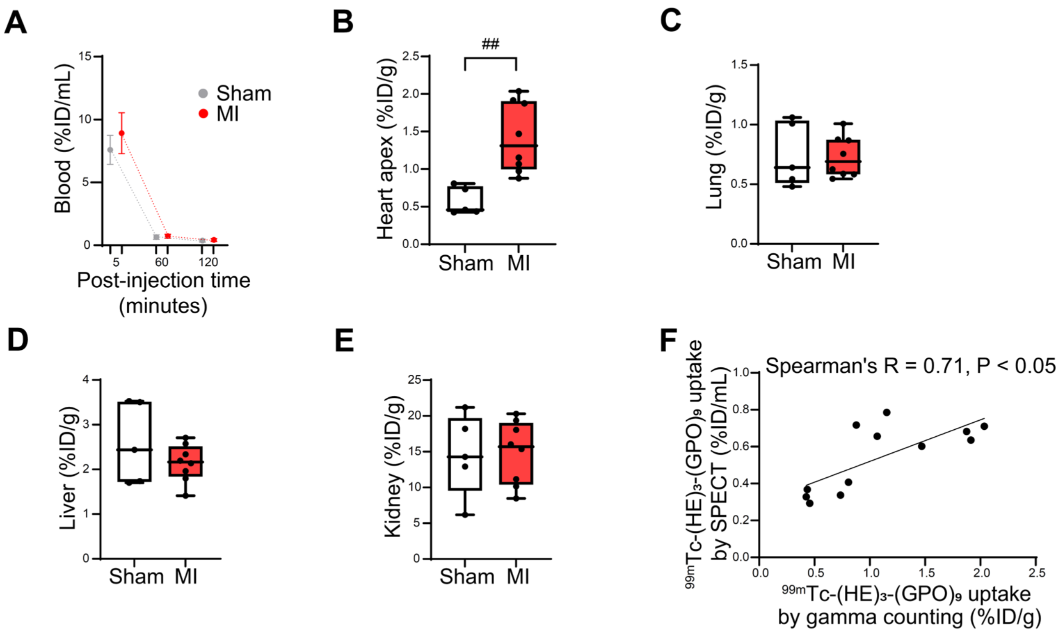


**Supplemental Figure 4: Biodistribution of $^{99m}$Tc-(HE)$_3$-(GPO)$_9$.** (**A**) Blood activity and tracer uptake in the heart apex, lungs, liver, and kidneys, measured with gamma counting, at 2 hours after tracer administration in 2-week post-MI and sham-operated animals. (**B**) Correlation between $^{99m}$Tc-(HE)$_3$-(GPO)$_9$ signal in the infarct zone and the anterolateral wall of sham-operated mice on in vivo SPECT/CT images and tracer uptake in the heart apex measured by ex vivo apical gamma counting. ID: injected dose, ##: P < 0.01, Mann-Whitney *U* test.

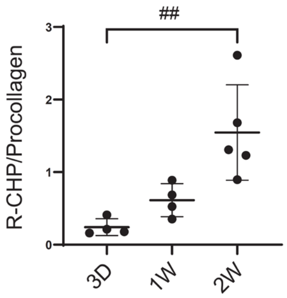


**Supplemental Figure 5: Collagen production and denaturation.** Quantification of the infarct zone denatured collagen (detected by R-CHP staining)-to-procollagen expression at different times after myocardial infarction. ##: P < 0.01, Kruskal-Wallis test.
